# Quantitative Relationship between Intracellular Metabolic Responses against Nutrient Conditions and Metabolic Inhibitions

**DOI:** 10.1101/2022.10.26.513808

**Authors:** Jumpei F Yamagishi, Tetsuhiro S Hatakeyama

## Abstract

Many previous studies have attempted to predict the metabolic states of cells assuming metabolic regulation is optimized through (sometimes artificial) evolution for some objective, e.g., growth rate or production of some metabolites. Conventional approaches, however, require identifying the microscopic details of individual metabolic reactions and the objective functions of cells, and their predictions sensitively depend on such details. In this study, we focus on the responses of metabolic systems to environmental perturbations, rather than their metabolic states themselves, and theoretically demonstrate a universal property of the responses independent of the systems’ details. With the help of a microeconomic theory, we show a simple relationship between intracellular metabolic responses against nutrient abundance and metabolic inhibition due to manipulation such as drug administration: these two experimentally measurable quantities show a proportional relationship with a negative coefficient. This quantitative relationship should hold in arbitrary metabolic systems as long as the law of mass conservation holds and cells are optimized for some objectives, but the true objective functions need not be known. Through numerical calculations using large-scale metabolic networks such as the *E. coli* core model, we confirmed that the relationship is valid from abstract to detailed models. Because the relationship provides quantitative predictions regarding metabolic responses without prior knowledge of systems, our findings have implications for experimental applications in microbiology, systems biology, metabolic engineering, and medicine, particularly for unexplored organisms or cells.

---

Prediction of cellular metabolic states is a central problem in biology. In particular, predicting the responses of metabolic systems against environmental variations or experimental operations is essential for controlling cellular states. Manipulating metabolic systems to the desired state is important in both life sciences and in applications such as matter production in metabolic engineering [1] and the development of drugs targeting cellular metabolism [2–4]. However, predicting the responses of metabolic systems remains a challenging task.

Previous studies have mainly attempted to predict the metabolic responses by predicting the metabolic states before and after perturbations; however, there are several problems. The conventional approach requires building an *ad hoc* model for each specific metabolic system. In systems biology, constraint-based modeling (CBM) has often been used to predict the cellular metabolic states [5, 6]. In this method, the intracellular metabolic state is predicted by solving an optimization problem of models of metabolic systems, including a detailed description of each metabolic reaction. To construct the optimization problem, metabolic systems of cells are assumed to be optimized through evolution for some objectives [5–7], e.g., maximization of the growth rate for reproducing cells such as cancer cells and microbes and maximization of the production of some molecules in metabolically engineered cells. The assumption is acceptable. Indeed, metabolic systems of reproducing cells exhibit certain universal phenomena across various species, and they can be explained as a result of optimization under physicochemical constraints [8]. Furthermore, metabolically engineered cells are sometimes strongly selected through artificial evolution [9]. However, knowing the true objective function of cells, which is essential to make a model for CBM, remains nearly impossible. Besides, even with remarkable progress in omics research, fully reconstructing metabolic network models for each individual species or cell of interest is still a challenge. In addition, the numerical predictions are sensitive to the details of the concerned constraints and the objective functions selected [10–13]. Therefore, predicting the metabolic state of arbitrary systems is difficult using the conventional approach, and we require new methods independent of the details of metabolic systems.

Instead of metabolic states themselves, here, we focus on the responses of metabolic systems to perturbations. At first glance, such prediction is seemingly more difficult than predicting the cellular metabolic states because it seems to require information not only on the steady states but also on their neighborhoods. However, from another perspective, to predict only the metabolic responses, we may need to understand the structure of only a limited part of the state space of feasible metabolic states. In contrast, we must seek the whole space to predict the metabolic states themselves. If optimization through evolution and some physicochemical features unique to metabolic systems constrain the behavior in the state space, there might be universal features in the responses of metabolic systems to perturbations, independent of system details.

In the present paper, we demonstrate a universal property of intracellular metabolic responses in the optimized metabolic regulation, using a microeconomic theory [14–16]. By introducing a microeconomics-inspired formulation of metabolic systems, we can take advantage of tools and ideas from microeconomics such as the Slutsky equation that describes how consumer demands change in response to the income and price. We thereby derive quantitative relations between the metabolic responses against nutrient abundance and those against metabolic inhibitions, such as addition of metabolic inhibitors and leakage of intermediate metabolites; the former is easy to measure in experiments while the latter may be not. The relations universally hold independent of the details of metabolic systems as long as the law of mass conservation holds. Our finding can be applied to any metabolic systems and will provide quantitative predictions on the intracellular metabolic responses without detailed prior knowledge of microscopic molecular mechanisms and cellular objective functions.

## RESULTS

### Microeconomic formulation of optimized metabolic regulation

We first provide a microeconomic formulation of metabolic regulation, which is equivalent to linear programming (LP) problems in CBM (Fig. 1; see also Appendix A in Supplemental Material for details).

**FIG. 1.**
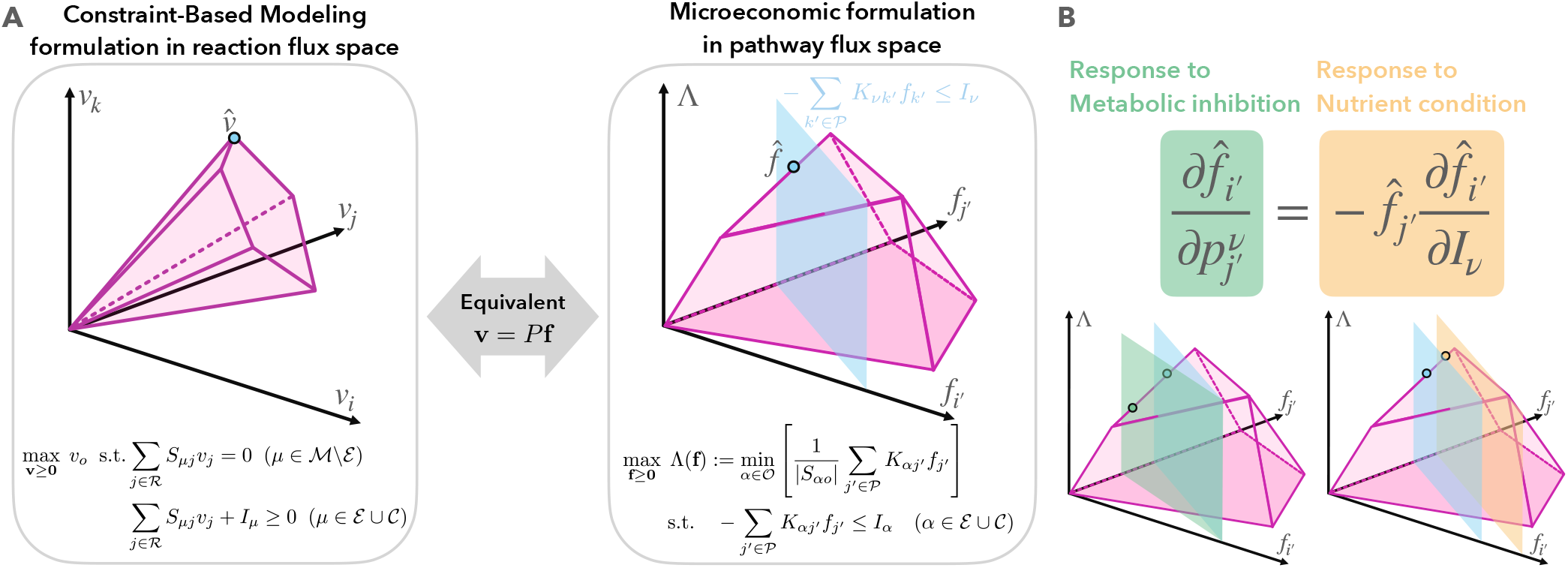
Schematic illustration of the metabolic CBM formulation and a microeconomic formulation. (A) (left) The metabolic CBM formulation with reaction fluxes **v** as variables. The solution space (convex set of possible allocations), called flux cone, in **v**-space is shown in pink. (right) Optimization of an objective flux Λ with pathway fluxes **f** as variables. The pink area in **f**-plane (bottom surface) represents the solution space, whereas the blue plane vertical to **f**-plane is the budget constraint for a component *ν*. The points 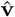 and 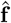 represent the optimized fluxes of reactions and pathways, respectively. Given that **v** = *P***f** holds with pathway matrix *P*, both representations and optimization problems are equivalent (see Appendix A in Supplemental Material for details). (B) The relation between the metabolic responses to changes in nutrient conditions (yellow) and those to metabolic inhibitions (green) [Eq. (3)].

We denote the set of all chemical species (metabolites) and that of all constraints by 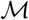 and 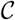, respectively. 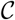 can reflect every type of constraint such as the allocation of proteins [17, 18], intracellular space [19], membrane surfaces [20], and Gibbs energy dissipation [13] as well as the bounds of reaction fluxes.

In the microeconomic formulation, variables to be optimized are the fluxes of metabolic pathways, whereas they are fluxes of reactions in usual CBM approaches; a metabolic pathway is a linked series of reactions and thus comprises multiple reactions. The sets of reactions and pathways are denoted by 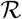 and 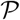, respectively. Let us then consider two stoichiometry matrices for reactions and pathways, *S* and *K*, respectively (see also Table I). If *α* is a chemical species 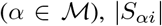 represents the number of units of species *α* produced for *S_αi_* > 0 and consumed for *S_αi_* < 0 in reaction *i*, whereas if *α* is a constraint 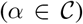, *S_αi_* is usually negative and |*S_αi_*| represents the number of units of constraint *α* required for reaction *i*. The other stoichiometry matrix *K* for metabolic pathways 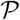 is also defined similarly. Throughout the paper, we use indices with primes such as *i* to denote pathways and those without primes such as *i* to denote reactions, and |*S_αi_*| and |*K_αi′_*| are called input (output) stoichiometry coefficients of reaction *i* and pathway *i′*, respectively, if *S_αi_* and *K_αi′_*, are negative (positive).

**TABLE I.** Description of the symbols in the main text

| Symbol | Description |
| --- | --- |
| $\mathcal{M}, \mathcal{C}, \mathcal{R}, \mathcal{P}$ | Set of chemical species / constraints / reactions / pathways |
| $\mathcal{E}, \mathcal{O}$ | Set of exchangeable species ($\mathcal{E} \subset \mathcal{M}$) / objective components ($\mathcal{O} \subset \mathcal{M} \cup \mathcal{C}$) |
| $S, K$ | Stoichiometry matrix for reactions / pathways ($S_{\alpha i}, K_{\alpha i'} \in \mathbb{R}; \alpha \in \mathcal{M} \cup \mathcal{C}, i \in \mathcal{R}, i' \in \mathcal{P}$) |
| $v_i, f_{i'}$ | Non-negative flux of reaction $i$ ($\in \mathcal{R}$) / pathway $i'$ ($\in \mathcal{P}$) |
| $q_i^\nu, p_{i'}^\nu$ | Metabolic price of reaction $i$ / pathway $i'$ in terms of $\nu$ ($\in \mathcal{M} \cup \mathcal{C}$) |
| $I_\alpha$ | Maximal intake of species $\alpha$ ($\in \mathcal{M}$) or total capacity for constraint $\alpha$ ($\in \mathcal{C}$) |
| $\Lambda$ | Flux of objective reaction $o$ ($\in \mathcal{R}$) |

Cells are assumed to maximize the flux of some objective reaction 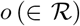 such as biomass synthesis in reproducing cells and ethanol or ATP synthesis in metabolically engineered cells. Objective reaction *o* requires objective components 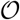, where *S_αo_* for each objective component 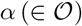 is negative because they are consumed in the reaction *o*. Because the reactants of a reaction cannot be compensated for each other due to the law of mass conservation, the flux is limited by the minimum available amount of objective components 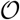 [16, 21, 22]. Therefore, the flux of objective reaction *o*, i.e., the objective function, is given as follows:

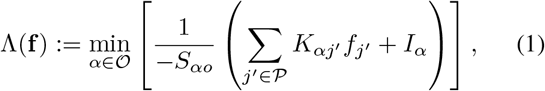

where 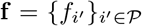 represents the fluxes of metabolic pathways. The arguments of the min function in Eq. (1) represent biologically different quantities: if *α* is a species 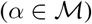, *I_α_* is the intake flux of species *α* and ∑*_j′_ K_αj′_ f_j′_* represents the total production rate of *α*, while if *α* is a constraint 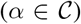, *I_α_* is the total capacity for constraint *α* and ∑*_j′_ K_αj′_ f_j′_* is the amount of *α* that can be allocated to the objective reaction.

Because the total consumption of exchangeable species 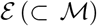 that can be transported through the cellular membrane cannot exceed their intakes, the available pathway flux **f** is subject to the following constraints in maximization of Λ(**f**):

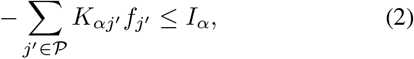

where *α* is in 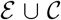 but not in 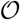. If species *α* is produced by objective reaction *o*, the intake effectively increases and *S_αo_Λ* is thus added to the right-hand side of Eq. (2), although this is not the case for most species. Therefore, the optimized solution f is determined as a function of *K* and **I**.

This optimization problem (1-2) can be interpreted as a microeconomic problem in the theory of consumer choice [14–16], considering Λ(**f**) as the utility function. By focusing on an arbitrary component 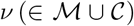, one of inequalities (2) serves as the budget constraint for *ν* if *K_νj′_*, ≤ 0 for all pathways *j′*, while the remaining inequalities in Eq. (2) then determine the solution space (Fig. 1A): for example, if we choose glucose as *ν*, the corresponding inequality in Eq. (2) represents carbon allocation. Here, the maximal intake *I_ν_* of *ν* corresponds to the income, and the input stoichiometry co-efficient for each pathway, 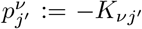, serves as the price of pathway *j′* in terms of *ν*.

### Relation between responses of pathway fluxes to nutrient abundance and metabolic inhibition

Because Eqs. (1-2) can be interpreted as an optimization problem in the theory of consumer choice, we can apply and generalize the Slutsky equation in microeconomics, which shows the relationship between changes in the optimized demands for goods in response to the income and price. In metabolism, this relation corresponds to that between the responses of optimal pathway fluxes 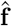 (see Appendix C.1 in Supplemental Material for derivation):

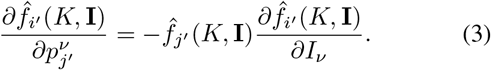

The right-hand side represents the responses of pathway *i′* against increases in *I_ν_*, whereas the left-hand side represents those against metabolic inhibitions in pathway *j′* because the metabolic price 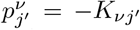, quantifies the inefficiency of conversion from substrate *ν* to end-products in pathway *j′* [16].

Eq. (3) is based on only the law of mass conservation: chemical reaction systems including metabolic systems necessarily have non-substitutability of reactants, i.e., the reactants of a reaction cannot be compensated for each other. Because the law of mass conservation stands in every chemical reaction, the relation (3) on the two measurable quantities must hold in arbitrary metabolic systems as long as their metabolic regulation is optimized for a certain objective. In particular, the case *i′* = *j′* will be useful: it follows that the measurement of the responses of a pathway flux to changes in the nutrient environment provides quantitative predictions of the pathway’s responses to metabolic inhibition or activation, and vice versa.

To confirm the validity of Eq. (3), we numerically solved the optimization problems (1-2) with pathway fluxes **f** as variables using the *E.coli* core model [5, 24] and randomly chosen stoichiometry coefficients for a single constraint (Fig. 2A; see also Appendix B in Supplemental Material for details). In this numerical calculation, metabolic pathways from exchangeable species to objective components are chosen as linear combinations of extreme pathways or elementary flux modes [25], i.e., extreme rays of the flux cone, for stoichiometry without objective reaction *o* (Fig. 2B), although the above arguments do not depend on the specific choices of metabolic pathways (see Appendix B in Supplemental Material for details). As shown in Fig. 2A, the quantitative relation (3) on the metabolic responses is indeed satisfied. Notably, the relation (3) itself invariably holds regardless of the number and type of constraint(s) 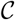 (see also Fig. S1 in Supplemental Material), whereas the metabolic states themselves can change sensitively depending on the concerned constraints and environmental conditions.

**FIG. 2.**
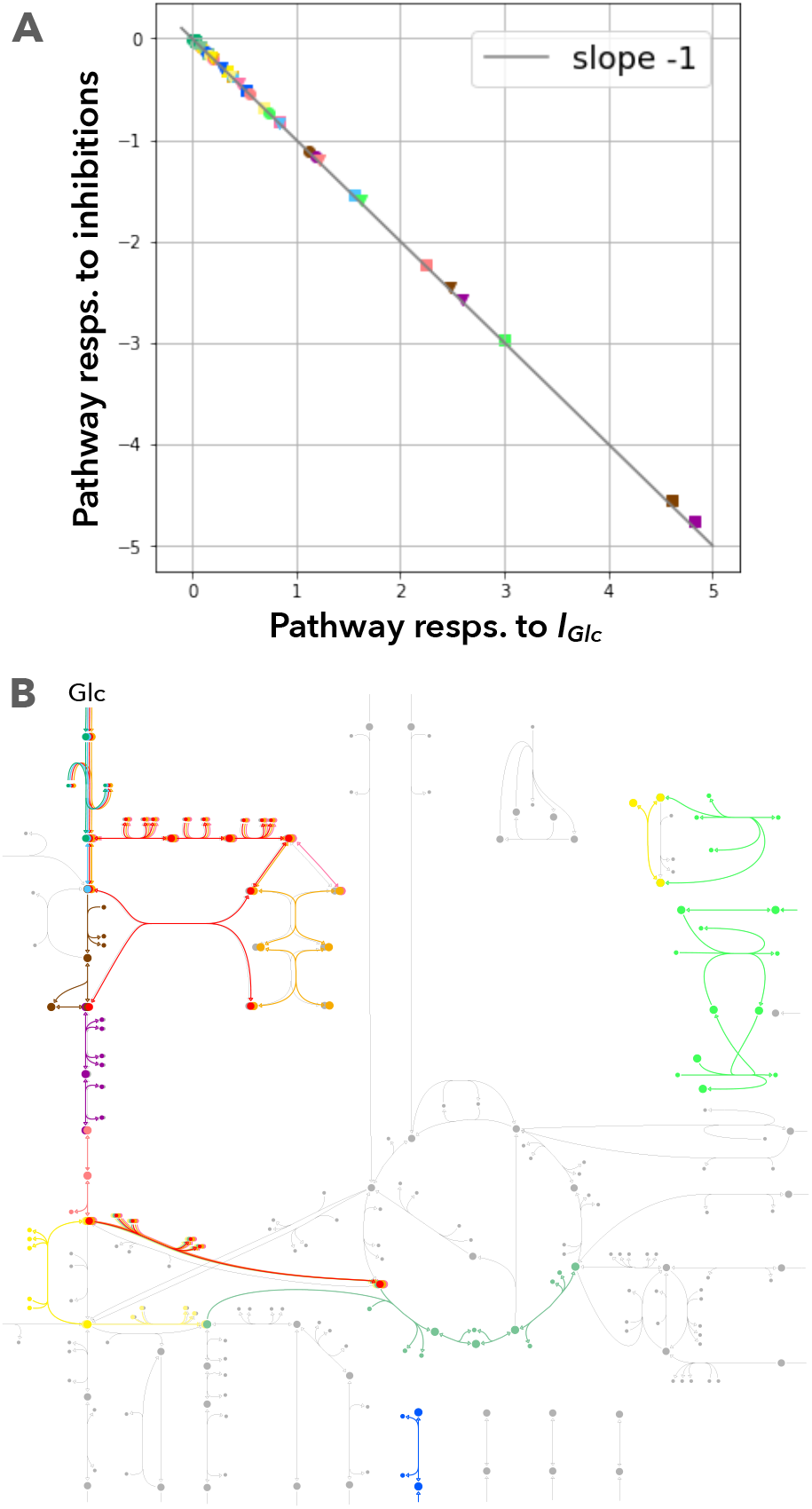
Metabolic responses of the optimized pathway fluxes 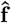. (A) Responses to metabolic inhibitions, 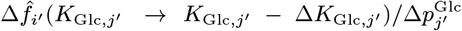, are plotted against the nutrient responses, 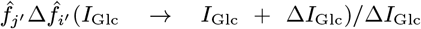. All different shapes and colors of markers represent different *i′* and *j′*, respectively. *I*_Glc_ = 5 [mmol/gDW/h], Δ*I*_clc_ = 0.1[mmol/gDW/h], Δ*K*_Glc,j′_/*K*_Glc,j′_ = 0.005. (B) 13 active extreme pathways, computed using efmtool [23], are shown using different colors that correspond to the markers for manipulated pathways *j′* in panel (A). The whole metabolic network of the *E. coli* core model is shown in light-gray.

### Relation between responses of reaction fluxes

Although Eq. (3) generally holds for every pathway, it may be experimentally easier to manipulate a single metabolic reaction. Manipulation of a single reaction can affect multiple pathways because multiple pathways and species are often tangled via a common reaction in metabolic networks. Thus, we should consider contributions of multiple pathways. The simplest way for this is to sum up Eq. (3) for all the pathways that include the perturbed reaction *i*. However, to precisely conduct this summation, we need to know the whole stoichiometry matrix or metabolic network. Hence, another relation closed only for the reaction fluxes **v** is required for application without the need to know the details of the metabolic systems.

To derive such a relation, we consider effective changes in the stoichiometry coefficients *S_αi_* for reaction *i* as metabolic inhibitions: e.g., inhibition of enzymes, administration of metabolite analogues, leakage of metabolites, and inefficiency in allocation of some resource. We then obtain an equality on the optimized reaction fluxes 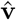, formally similar to Eq. (3) (see Appendix C.2 in Supplemental Material for derivation):

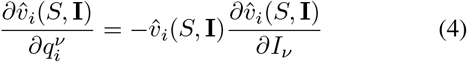

by defining the metabolic price 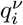 of reaction *i* in terms of *ν* as a function of *S*, instead of the metabolic price 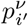 of pathway *i′* as a function of *K*,

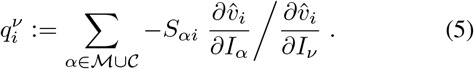

The coefficient 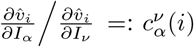 quantifies the number of units of component *ν* that can compensate for one unit of *α* in reaction *i* and is experimentally measurable. For example, if *ν* is glucose and *α* is another metabolite such as an amino acid, 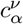 indicates how many units of glucose are required for the compensation for one unit of the amino acid, similar to the “glucose cost” in previous studies [26].

For the calculation of Eq. (4), changes in the metabolic price 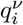 need to be calculated rather than the metabolic price itself. The price changes depend on the type of manipulations we focus on: (I) manipulations leading to the loss of a single component and (II) those leading to the loss of multiple components.

#### (I) Metabolic inhibition of a single component

If experimental manipulation causes the loss of a single component 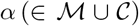 in reaction *i, S_αi_* effectively changes only for that *α* (Fig. 3A). In such a case, the metabolic price change is just given by 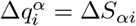.

**FIG. 3.**
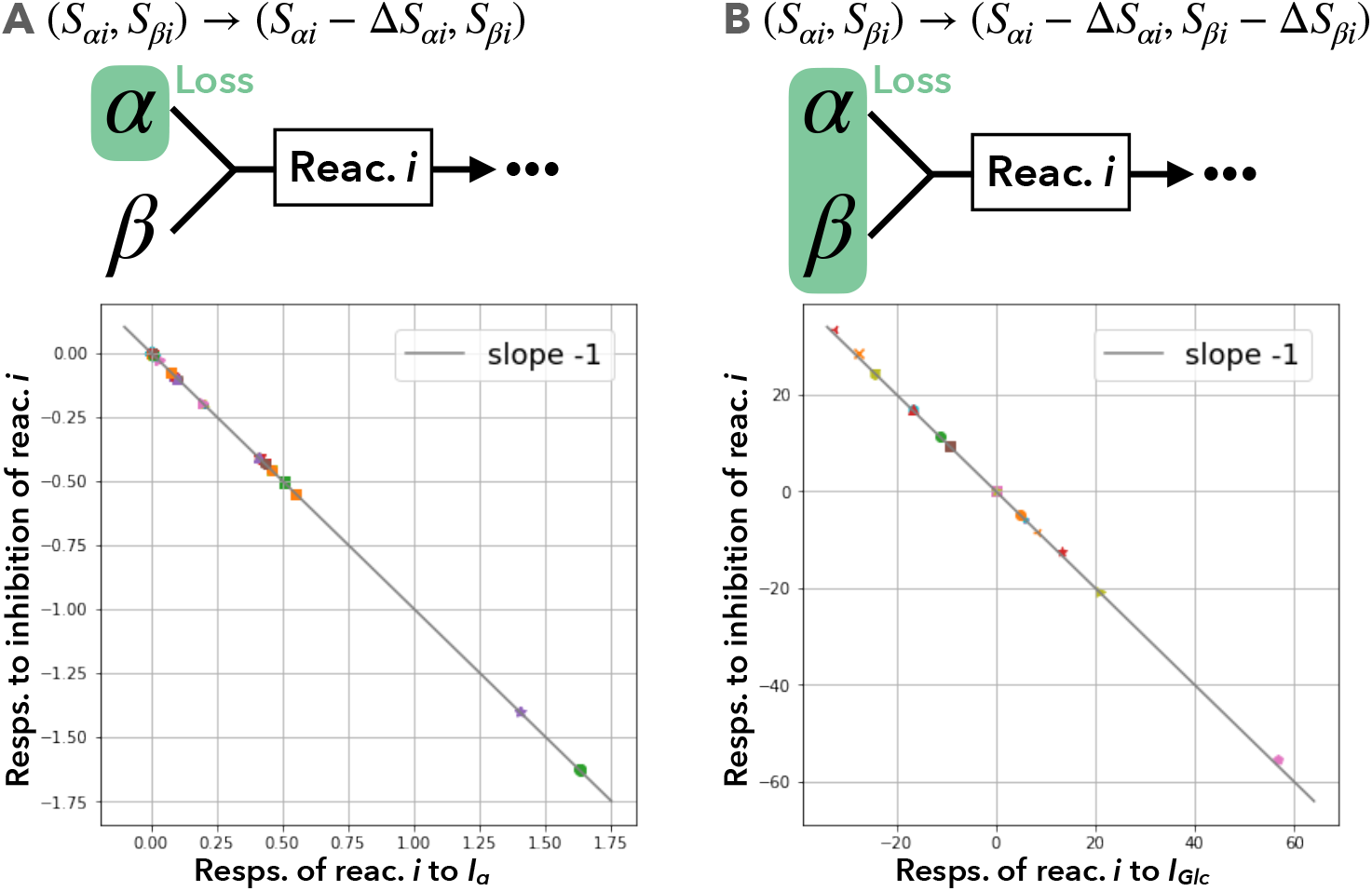
Metabolic responses of the optimized reaction fluxes 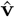 against metabolic inhibitions of reaction *i*. As the simplest example, we illustrate a two-body reaction for *α* and *β* in the upper panels. (A) Manipulation that leads to the loss of only a single species (*a* in top panel): case (I). The horizontal axis shows the responses to the available amount of a constraint 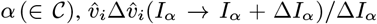, whereas the vertical axis does those to metabolic inhibitions, 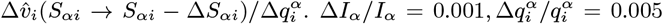. (B) Manipulation that leads to the loss of multiple species (*α* and *β* in top panel): case (II). Responses of the optimized reaction flux 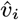 to effective changes in the input stoichiometry coefficients 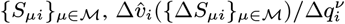, are plotted against those to intake changes, 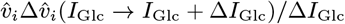. *I*_Glc_ = 8.05 [mmol/gDW/h], Δ*I*_Glc_ = 0.0025 [mmol/gDW/h], 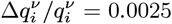. Each marker corresponds to a different reaction *i*.

An example of such experimental manipulations is the administration of an analogue to a reactant of a multi-body reaction: if *α* and *β* react (see Fig. 3A), the metabolic analogue of *β* can produce incorrect metabolite(s) with *α*, leading to the loss of *α*, and thus, reaction *i* requires more *α* to produce the same amount of products, causing effective increases in the input stoichiometry coefficients |*S_αi_*|. Another example is the changes in the total capacity and effective stoichiometry for a constraint: for example, the mitochondrial volume capacity will work as such a constraint and can be genetically manipulated [27–29]. As is numerically confirmed in Fig. 3A, Eq. (4) is indeed satisfied even in such a case.

#### (II) Metabolic inhibition of multiple components

If metabolic inhibition of multiple reactant species of a reaction *i* is simultaneously caused, the stoichiometry coefficients *S_μi_* for multiple reactants *μ* of reaction *i* will effectively change (Fig. 3B). Accordingly, the reaction price changes by 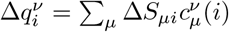.

In experiments, such cases would correspond to inhibition of enzymes, leakage of the intermediate complex of reaction *i*, and so forth.

Even in such cases, the quantitative relation (4) is satisfied, as is confirmed by numerically calculating the price changes of reaction *j* defined in Eq. (5) (Fig. 3B). Although the precise calculation of the coefficients 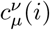 requires information regarding not only the responses of 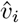, to *I_ν_* but also those to *I_μ_*, they can be approximated in ways easier and independent of reaction *i*. For example, 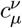 should be the ratios of the carbon numbers of species *μ* and *ν* under extreme situations in which only the carbon sources limit the objective reaction; alternatively, the simplest approximation could be just taking 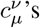 as unities. Even with these approximated values of 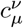, the relation (4) is still valid approximately (see Fig. S1 in Supplemental Material).

##### Independence of the relation between the metabolic responses on cellular objective functions

Finally, our above argument does not depend on specific choices of the objective reaction *o*, whereas we have utilized the biomass synthesis reaction as *o* in Figs. 2 and 3. To highlight the independence of the relation (4) from cellular objective functions, we conducted another numerical calculation (Fig. S2 in Supplemental Material). The relation (4) is indeed satisfied even when the objective reaction *o* is set as a reaction for matter production, such as ethanol or ATP synthesis, which is often considered as the objective for metabolically engineered cells [1, 30, 31].

## DISCUSSION

In the present study, we showed that the metabolic responses against resource availability and those against metabolic inhibitions are negatively proportional. The quantitative relations we found should be universally satisfied with arbitrary reaction networks, constraints, and objective functions of cells. In particular, although the predicted optimal metabolic states can drastically depend on the assumed objective function, the relations on the responses should be always satisfied independent of it (Fig. S2 in Supplemental Material). Although we can never know the true objective function of cells, we can still predict the metabolic responses.

The independence of the cellular objective functions could be derived from the microeconomic formulation for metabolic regulation, equivalent to CBM, and application of the Slutsky equation in economics. Notably, the Slutsky equation in economics generally requires detailed information regarding the objective functions (i.e., utility functions in economics) because it includes a term for the so-called substitution effect that quantifies the substitutability of goods and depends on the objective functions (see also Sec S3.A in Supplemental Material). However, the term disappears in the case of metabolic systems because of the law of mass conservation. Thus, the relation can be easily calculated and will be experimentally testable without prior knowledge of cellular objective functions.

Because our finding is valid regardless of how abstract the concerned model is, from coarse-grained toy models to genome-scale metabolic networks, the relation we found would be important both for quantitative predictions and for discovering qualitatively novel phenomena. The Warburg effect is a prominent example of the latter, which has sometimes been explained based on coarse-grained models comprising two metabolic pathways for respiration and fermentation [8, 32]. In the Warburg effect, if the amount of the carbon source taken up by a cell increases, the cell decreases the flux of the respiration pathway [33]. From the relation (3) between responses, we can immediately predict that an increase in the metabolic price of respiration (e.g., administration of uncouplers of respiration [34]) will counterintuitively increase the respiration flux. Indeed, in the previous studies, such an increase in the respiration flux was observed in a coarsegrained model, which was termed the drug-induced reverse Warburg effect [16]. This phenomenon was also observed in several published experiments [34–37]. In economics, the phenomenon in which individuals consume goods more when their price increases is called Giffen behavior. Thus, the drug-induced reverse Warburg effect is a prominent example of Giffen behavior in metabolism, and numerous other seemingly counterintuitive metabolic behaviors might be understood from the viewpoint of Giffen behavior.

In experimental application, the intake or total capacity can be altered by shifts in environmental conditions, genetic manipulations, and so forth, whereas changes in the metabolic price can be also implemented in various ways: e.g., administration of a metabolite analogue, leakage of a metabolite, and inhibition of some enzyme that will lead to accumulation of the reactants and possibly promote their excretion or conversion to other chemicals. They cause a loss of reactants, and thus, the corresponding reaction(s) require more metabolites to produce the same amount of products.

The relations (3-4) allow us to predict the responses of a reaction or pathway flux of interest to metabolic inhibitions only by measuring its fluxes in several different nutrient conditions, and vice versa. Remarkably, the predictions do not require detailed information regarding the concerned intracellular reaction networks, and they are valid even when the precise estimation of effective changes in the stoichiometry coefficients is difficult, at least qualitatively (Fig. S2 in Supplemental Material). Therefore, they will be useful as a quantitative and qualitative guideline to operate the metabolic states toward the desirable directions in various fields such as microbiology, metabolic engineering, and drug development.

## Appendix A: Equivalence between constraint-based modeling (CBM) and microeconomic formulations

### 1. CBM formulation: optimization problems with reaction fluxes v as variables

In the framework of CBM in systems biology, intracellular metabolic regulation is formulated as linear programming (LP) problems in which the variables are the fluxes **v** of reactions. As discussed below, LP problems in CBM are generally equivalent to optimization problems in the microeconomic theory of consumer choice.

First, when each reversible reaction is broken down into two irreversible reactions (i.e., its forward and backward components), a non-negative 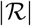-dimensional vector 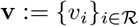 represents the fluxes of all reactions.

By assuming the stationarity of the intracellular concentrations of non-exchangeable species, the general formulation of CBM [5, 6] is given as follows:

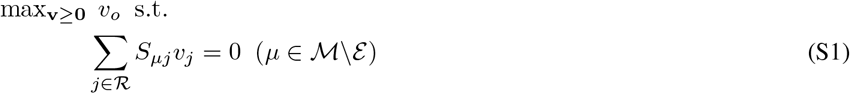

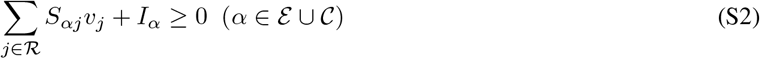

Because 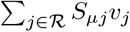 is equal to the excess production of species *μ*, Eq. (S1) thus represents that the production and degradation of internal metabolites must be balanced. With respect to Eq. (S2), if *α* is a species 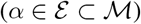, the corresponding inequality, 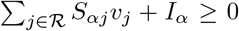, represents that the exchangeable species *α* with intake *I_μ_ >* 0 are taken in and species *μ* with efflux *I_μ_* < 0 are (forcibly) leaked or degraded; in contrast, if *α* is a constraint 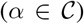, it represents one of the other constraints, e.g., allocation of some limited resource such as proteins [17, 18], intracellular space [8, 19], membrane surface [20], and Gibbs energy dissipation [13], where *S_αi_* for 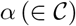 is typically negative or zero but can also be positive. Note that C can also include other constraints like the upper and lower bounds of the flux of reaction *i*.

### 2. Optimization problems with pathway fluxes f as variables

The usual CBM formulation (S1-S2) with reaction fluxes **v** as variables is equivalent to another LP problem with pathway fluxes **f** as variables (see also Fig. 1 in the main text).

Here, a metabolic pathway is a linked sequence of reactions. Pathway fluxes **f** are related to reaction fluxes as **v** = *P**f***, with a *pathway matrix* 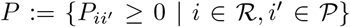 which represents that pathway *i′* comprises *P_ii″_* units of reaction *i*. The stoichiometry matrix *K* for pathways expresses the metabolic pathways in “species space” and is related to the stoichiometry matrix *S* for reactions with *K* = *SP*.

When *P* is taken as (linear combinations of) elementary flux modes (EFMs) (matrix *K*:= *SP* is then related to so-called elementary conversion modes [38]), Eq. (S1) is immediately satisfied. Then, the LP problem (S1-S2) for CBM can be rewritten into another LP problem with pathway fluxes **f** as variables:

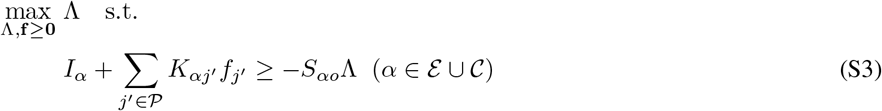

where the components required for objective reaction 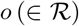 are termed as objective components 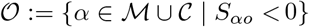.

### 3. Derivation of the microeconomic formulation

The min function for the objective function, Λ:= min(*A, B*), can be replaced by constraints of Λ ≤ *A* and Λ ≤ *B*, and vice versa. Therefore, the optimization problem (S3) is equivalent to the optimization of the Leontief utility function [Eq. (1) in the main text]—the minimum of multiple “complementary” objectives—under the constraints of Eq. (2) in the main text. In contrast, the microeconomic formulation of optimization problems [Eqs. (1-2) in the main text] can be converted to LP problems in the form of Eq. (S3), and thus, they are equivalent.

Throughout the numerical simulations in the main text, the intakes are set as *I*_O_2__ = 22.35 [mmol/gDW/h], *I*_Phos_ = 3.17 [mmol/gDW/h].

## Appendix B: Details of numerical experiments

In our numerical simulations, stoichiometric coefficients *S_αi_* for species 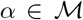 are given by the *E.coli* core model [24]. *S_αi_*’s for constraints 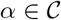 are randomly chosen from an interval [-1, 0] to show that the results do not depend on such details; here, as a natural assumption, we set *S_αi_’s* for the reactions corresponding to a reversible reaction as identical and those for the transport reactions via diffusion as zero.

In most numerical simulations, the objective reaction *o* is chosen as the biomass synthesis reaction, whereas our results do not depend on the choice of the objective reactions: e.g., synthesis reaction of ethanol or ATP (see also Fig. S2).

We plotted only reactions with continuous price responses in Figs. 2 and 3 in the main text and Figs. S1 and S2. In our numerical calculations, the response to price change is considered to be noncontinuous if decreasing Δ*p* changes the reaction flux discontinuously at Δ*p* = 0; or more easily, if ∥**v**(*p_i_* + Δ*p, I*) – **v**(*p, I*)∥/Δ*p* is much larger than ∥**v**(*p, I* + Δ*I*) – **v**(*p, I*)∥/Δ*I*.

### 1. On the uniqueness of the solutions

For simplicity, the arguments in the main text are based on the premise of the uniqueness of the solution. This assumption indeed holds true with suitable penalty terms explained below. Furthermore, it must be biologically natural because we considered the intracellular metabolic responses around a steady state here.

An LP can sometimes have multiple solutions. If an LP has multiple solutions because of the existence of ineffective inequalities, it is natural to define the unique solution as the solution with the least consumption rates of species: i.e., slightly modify the objective function as 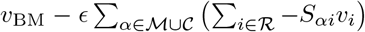 with small *ϵ* > 0 (here, note that 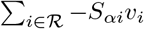 is equal to the consumption rate of *α*, while *I_α_* is its upper bound). In addition, for the pair of two irreversible reactions formed by separating each reversible reaction, the smaller of the two must be subtracted and set to zero for uniqueness of the solution by adding a penalty –*ϵ* ∑*_i_ ν_i_*. Throughout the numerical calculations in the main text, *ϵ* is basically set to ≲ 10^-5^, whereas the results do not depend on *ϵ* as long as *ϵ* is sufficiently small.

### 2. Leakage of intermediate species

To calculate the reaction price 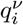 [Eq. (5) in the main text], we must calculate the responses of 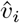 against changes in *I_μ_* for each reactant *μ* of reaction *i*. For calculating the responses to *I_μ_* of each *μ*, we assumed that species *μ* is exchangeable. Such changes do not alter the optimized solution 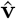 in the numerical calculations we conducted.

### 3. Choice of metabolic pathways in Fig. 2 in the main text

In this study, we consider (linear sums of) EFMs or extreme pathways for the stoichiometry without objective reaction *o* as the metabolic pathways. That is, we consider metabolic pathways from components *α* with influxes *I_α_ >* 0 to objective components 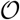.

## Appendix C: Derivation of Eqs. (3-5) in the main text

### 1. Derivation of Eq. (3): Slutsky equation in microeconomics

When given a utility function *u*(**x**), we define 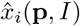, as the optimal demand for good *i* determined as a function of the price of the good **p** and income *I*. By defining *E*(**p**, *u*) as “the minimum income required to achieve a certain utility value *u*,” we can represent “the minimum demand for good *i* required to achieve a utility value *u*” as 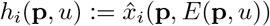.

Differentiating this function *h_i_*(**p**, *u*) with respect to *p_j_* yields

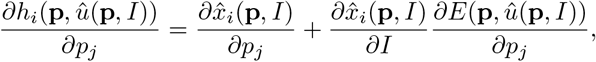

where 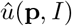 represents the maximum utility under a given price **p** and income *I*. Due to optimality, the last term 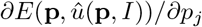 equals to the optimal demand 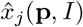.

Accordingly, we obtain the Slutsky equation that describes the response of demand 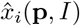 to changes in price *p_j_*:

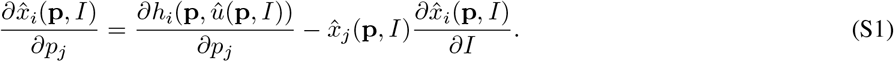

The first term 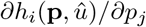 represents the substitution effect caused by relative changes in the price of each good; particularly, the “self-substitution effect” for *i = j* is always non-positive [14]. In contrast, the second term 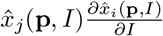 reflects the income effect, which can be either positive or negative. This represents the effect that an increase in the price of a good leads to an effective decrease in the income, which also changes the demand for goods.

The law of mass conservation in metabolism corresponds has the perfect complementarity in economics. Accordingly, the substitution effect is always null (at a kink) [15, 16]. Then, by noting that 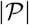 metabolic pathways correspond to goods in economics in our mapping between metabolism and microeconomics, we immediately obtain Eq. (3) in the main text from the Slutsky equation (S1).

### 2. Derivation of Eqs. (4-5) from Eq. (3) in the main text

Noting 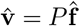 and multiplying Eq. (3) in the main text by pathway matrix *P* from the left, we immediately obtain an equality for the responses of reaction fluxes **v**:

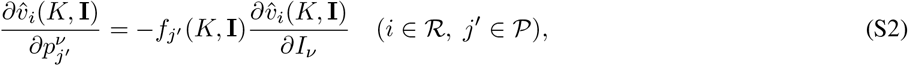

Assuming that a stoichiometry coefficient *S_μj_* of reaction *j* is effectively altered to *S_μj_* – Δ*S_μj_*, the *j′*-th column vector of matrix *K*:= *SP* changes as

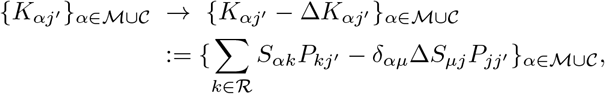

with Kronecker delta *δ_αμ_*. That is, the metabolic price of pathway *j′* for metabolite *μ* changes by 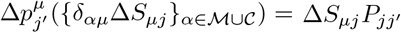.

Therefore, if multiple stoichiometry coefficients are simultaneously altered,

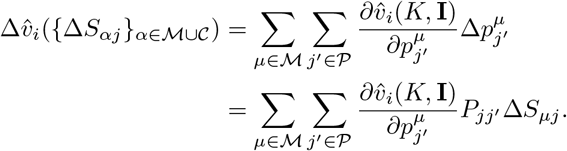

From Eq. (S2) and 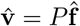,

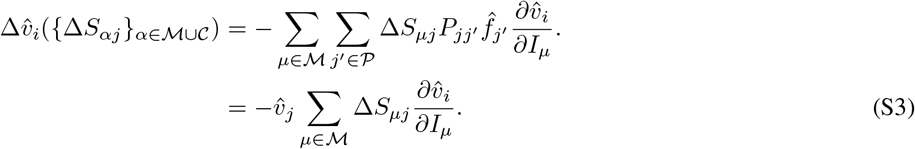

Then, by defining 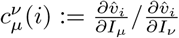 and

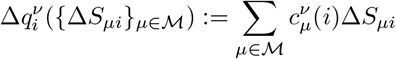

and applying it to Eq. (S3) in the case of *j* = *i*, we obtain

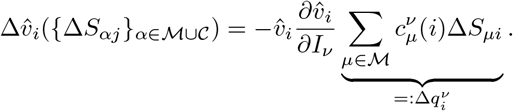

This is the derivation of Eqs. (4-5) in the main text.

**FIG. S1.**
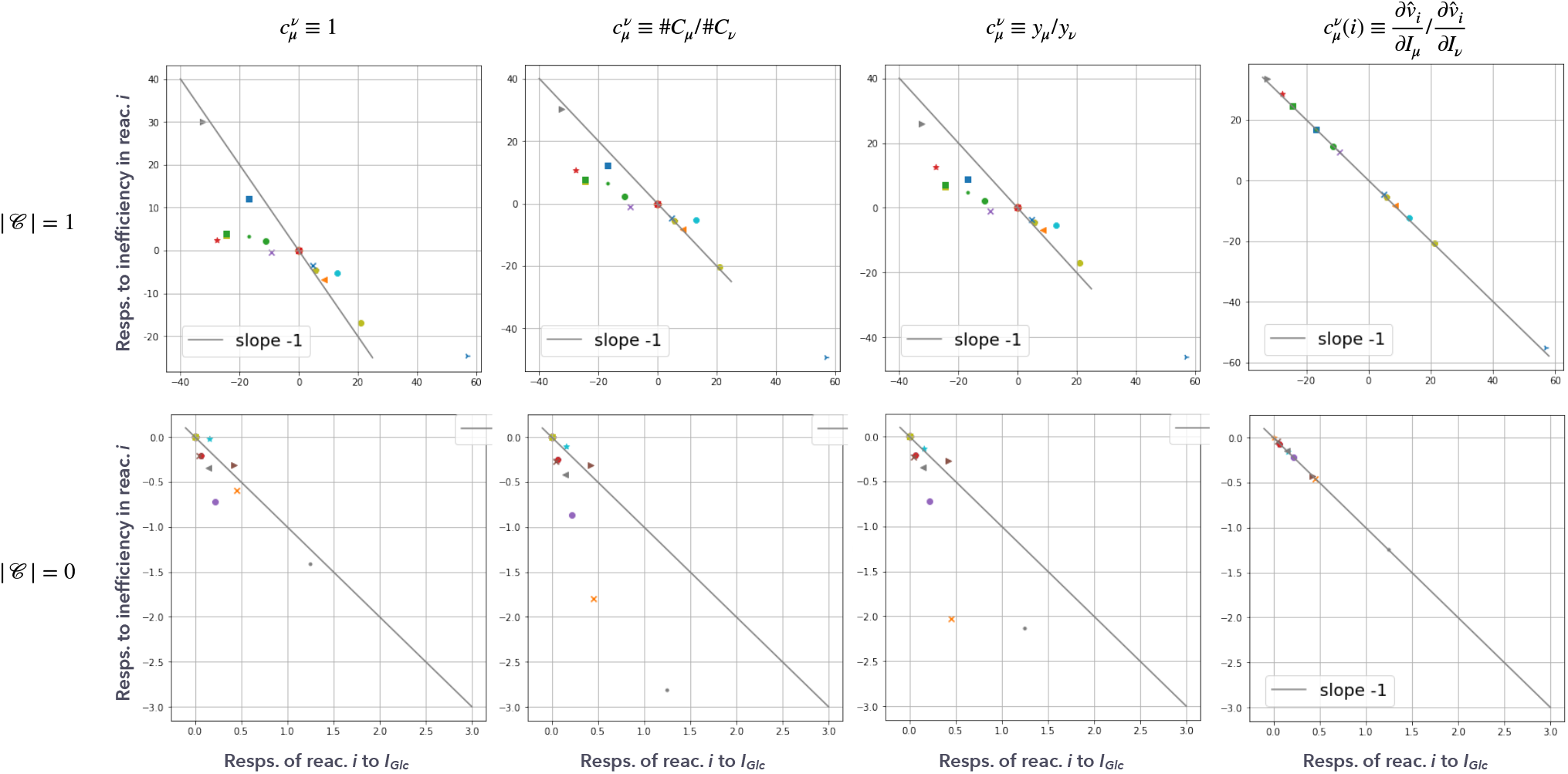
The metabolic responses with several approximated estimations of 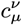 are plotted. (Top) With a constraint (i.e., the number of elements in 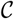 is 1). (Bottom) Without constraints (i.e., the number of elements in 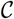 is 0). From the left, 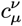 is approximated as 1 independent of *μ*, 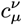 is approximated as the ratios of the carbon numbers of species *μ* and *ν*, 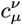 is approximated as 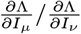 (that equals to the ratio of the shadow price [39] of species *μ* and *ν, y_μ_/y_ν_*), and Eq. (5) in the main text (without approximation).

**FIG. S2.**
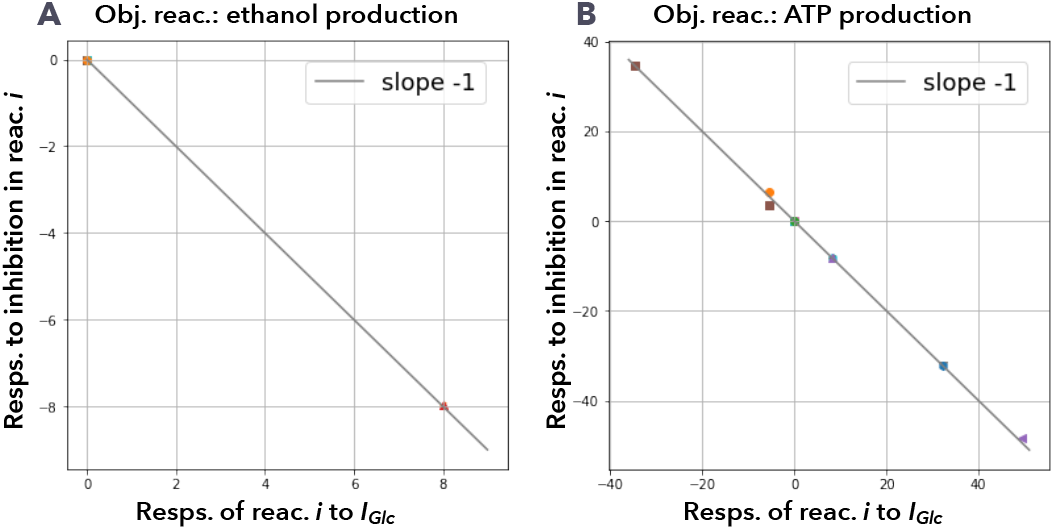
The metabolic responses with matter production as the objective reaction. (A) Ethanol production (acetaldehyde dehydrogenase; abbreviated as “ADHEr” in *E. coli* core model). (B) ATP production (ATP maintenance requirement; abbreviated as “ATPM” in *E. coli* core model).

